# Cellular reductase activity in uncultivated *Thiomargarita spp*. assayed using a redox-sensitive dye

**DOI:** 10.1101/165381

**Authors:** Jake V. Bailey, Beverly E. Flood, Elizabeth Ricci, Nathalie Delherbe

## Abstract

The largest known bacteria, *Thiomargarita spp.,* have yet to be isolated in pure culture, but their large size allows for individual cells to be followed in time course experiments, or to be individually sorted for ‘omics-based investigations. Here we report a novel application of a tetrazolium-based dye that measures the flux of reductase production from catabolic pathways to investigate the metabolic activity of individual cells of *Thiomargarita* spp. When coupled to microscopy, staining of the cells with a tetrazolium-formazan dye allows for metabolic responses in *Thiomargarita* spp. to be to be tracked in the absence of observable cell division. Additionally, the metabolic activity of *Thiomargarita* spp. cells can be differentiated from the metabolism of other microbes in specimens that contain adherent bacteria. The results of our redox-dye-based assay suggests that *Thiomargarita* is the most metabolically versatile under anoxic conditions where it appears to express cellular reductase activity in response to the electron donors succinate, acetate, citrate, formate, thiosulfate, H_2_, and H_2_S. Under hypoxic conditions, formazan staining results suggest the metabolism of succinate, and likely acetate, citrate, and H_2_S. Cells incubated under oxic conditions showed the weakest formazan staining response, and then only to H_2_S, citrate, and perhaps succinate. These results provide experimental validation of recent genomic studies of *Ca.* Thiomargarita nelsonii that suggest metabolic plasticity and mixotrophic metabolism. The cellular reductase response of bacteria attached to the exteriors of *Thiomargarita* also supports the possibility of trophic interactions between these largest of known bacteria and attached epibionts.

**IMPORTANCE:** The metabolic potentials of many microorganisms that cannot be grown in the laboratory are known only from genomic data. Genomes of *Thiomargarita* spp. suggest that these largest of known bacteria are mixotrophs, combining lithotrophic metabolisms with organic carbon degradation. Our use of a redox-sensitive tetrazolium dye to query the metabolism of these bacteria provides an independent line of evidence that corroborates the apparent metabolic plasticity of *Thiomargarita* observed in recently produced genomes. Finding new cultivation-independent means of testing genomic results is critical to testing genome-derived hypotheses on the metabolic potentials of uncultivated microorganisms.

## INTRODUCTION

Sulfide-oxidizing bacteria of the family Beggiatoaceae contain some of the largest known bacteria (1-5), with individual cells of the genus *Thiomargarita spp.* reaching millimetric diameters (3, 5, 6). Dense communities of these organisms on the seafloor comprise some of the most spatially-extensive microbial mat ecosystems on Earth (4, 7). The large sulfur bacteria are primarily chemolithotrophs or mixotrophs that live at interfaces between nitrate or oxygen, and hydrogen sulfide (8-12). These bacteria can have a substantial influence on the biogeochemical cycling of sulfur, nitrogen, phosphorus and carbon in diverse settings (11, 13-15). Despite their large size and biogeochemical significance, the physiologies and ecologies of these bacteria remain incompletely understood, in part because the large sulfur bacteria are not presently isolated in pure culture. At present, *Thiomargarita* spp. can only be maintained long-term in their natural sediments, not in axenic isolation or even as mixed communities in defined microbial media. Therefore approaches to studying *Thiomargarita*’s physiology have focused on culture-independent approaches. Schulz and deBeer used microsensors to measure O_2_ and sulfide gradients influenced by *Thiomargarita* metabolisms (16), and more recently, (meta)genomic and metatranscriptomic approaches have been used to investigate their genetic potential (17, 18) and response to environmental perturbations (19). Here we add to these findings using a novel approach to studying large sulfur bacteria that employs tetrazolium redox dyes.

Tetrazolium dyes are widely used for measuring the metabolic activity of cells, including both bacteria (20-22) and eukaryotes (23), growing on a variety of metabolic substrates. In their oxidized form, soluble tetrazolium salts generally result in a colorless or weakly-colored solution. Cellular reductases, such as NADH, reduce tetrazolium salts to an intensely-colored formazan product that can be observed qualitatively or measured quantitatively in a variety of colorimetric metabolic assays (24-26). These dyes can be used to measure catabolic metabolic activity even in the absence of observable cell division. Typically, tetrazolium dyes are applied to bacteria in culture and measured via color response of bulk media and culture in a microplate (27). We sought to apply a tetrazolium dye approach to investigate metabolism in *Thiomargarita* spp. However, *Thiomargarita spp.* are not in pure culture, and these large cells are covered with communities of smaller attached bacteria (12). Our initial attempts to analyze our custom tetrazolium microplate assays via spectroscopy failed to differentiate between the metabolism of *Thiomargarita* and its attached bacteria. We then employed a microscopy-based approach to image tetrazolium color change associated with individual *Thiomargarita* cells, so as to differentiate metabolic responses of *Thiomargarita* spp. from those of attached epibiont cells. Here we report on the cellular reductase response of *Thiomargarita* spp. cells to several organic and inorganic substrates under oxic, hypoxic, and anoxic conditions.

## MATERIALS AND METHODS

### Sample Collection

*Thiomargarita* spp. cells were collected from organic-rich sediments on the Atlantic shelf, near Walvis Bay, Namibia (23°00.009' 14°04.117') using a multi-corer on board the R/V *Mirabilis*. All cells used in the experiments were collected from a depth of 1-3 cm depth beneath the sediment/water interface. *Thiomargarita* sp. cells were stored in their host sediments with overlying core-top water in closed 50 ml centrifuge tubes at 4°C, and protected from direct light for three weeks.

### Incubation experiments with tetrazolium staining

Chains and clusters of *Thiomargarita* sp. cells were rinsed three times in 0.2 μm filtered artificial seawater before being added to an incubation medium in 96-well microplates. The purpose of these rinse steps was to remove loosely attached smaller bacteria from the exteriors of *Thiomargarita* cells and sheaths. These wash steps did not remove tightly-attached or sheath-embedded epibionts, as confirmed by microscopy and described below. One *Thiomargarita* cell, cell cluster, or cell chain, was added to each individual well in the 96-well plate. Each substrate treatment contained eight wells with *Thiomargarita* cells, as well as four control wells without *Thiomargarita* cells. Two of these control wells contained 20 μl of the first saltwater bath used to rinse the *Thiomargarita* cells, added to 180 μl of media, as a control that contains cells found loosely attached to *Thiomargarita*. Wells containing empty diatom frustules picked by pipette from the same samples as the *Thiomargarita* cells were also used as negative controls to monitor color change of xenic biological surfaces from the same environment. A third control type consisted of empty mucus sheaths that were produced by *Thiomargarita*, but which no longer contained *Thiomargarita* cells. These empty sheaths are common in *Thiomargarita*-rich sediments and can be readily identified by the shape of the chains that is preserved as void space in the sheath material.

The incubation medium was designed to maintain metabolizing *Thiomargarita* cells and to provide basic elemental constituents and nutrients based on similar base media recipes for phylogenetically-related taxa (28). The incubation medium included the following constituents (l^-1^): NaCl, 34 g; CaCl_2_, 0.112g; NH_4_NO_3_, 0.008g; KCl, 0.5g; MgSO_4_, 1.46; 20 mL 1M MOPS (3-(*N*-morpholino)propanesulfonic acid) buffer (pH adjusted to 7.8, final conc. 20mM); 1 ml 1000x potassium phosphate buffer (pH 7.6); 1 ml 1000x B_12_; 1 ml 1000x vitamin solution, 10 ml 100x trace element solution. The vitamin solution contained (l^-1^) 10 mg of riboflavin and 100 mg each of thiamine HCl, thiamine pyrophosphate, L-ascorbic acid, D-Ca-pantothenate, folic acid, biotin, lipoic acid, nicotinic acid, 4–aminobenzoic acid, pyridoxine HCl, thiotic acid, nicotinamide adenine dinucleotide, and inositol dissolved in 100 mL of a 10 mM KPO_4_ buffer (pH 7). The trace metal solution contained (l^-1^) 0.1 g FeCl_2_ • 2H_2_O, 0.03 g H_3_BO_3_, 0.1 g MnCl_2_, 0.1 g CoCl_2_ • 6H_2_O 1.5 g nitrilotriacetic acid, 0.002 g NiCl_2_ • 6H_2_O, 0.144 g ZnSO_4_ • 7H_2_O, 0.036 g NaMoO_4_, 0.025 g Na-vanadate, 0.010 g NaSeO_3_ and NaWO_4_ • 2H_2_O. NaHCO_3_ was added to each base medium for a final concentration of 3mM for the oxic and anoxic stocks, and 40 mM for the hypoxic treatments. The medium was sterilized by filtration through 0.22 μm membrane, and the final pH was 7.9-8.0.

Potential electron donors acted as experimental variables, and included H_2_, H_2_S, thiosulfate, succinate, acetate, citrate, and formate. All electron donors except for H_2_ and H_2_S were supplied at final concentrations of 1mM. H_2_ was supplied by shaking the plates in a Coy anaerobic chamber containing a 3% H_2_, 98% N_2_ atmosphere. H_2_S was supplied by the daily addition of 10 μl of the freshly-neutralized 4mM sodium sulfide delivered by syringe. The tetrazolium redox dye mix we used, which is known by the commercial name “Dye H” (Biolog Cat. No 74228) was added to the medium at a final 1x working strength just prior to cell incubation.

Microplates were placed on orbital shakers at 50 rpm. One microplate was maintained under benchtop atmospheric conditions, one microplate was placed in a Coy hypoxic chamber with 5% atmospheric level O_2_ and ∼5% total CO_2_, one microplate was placed in an H_2_-free anaerobic chamber (NextGen, Vacuum Atmospheres Company, Hawthorne, CA), and one microplate was placed in a Coy anaerobic chamber containing a gas mixture 97% N_2_, 3% H_2_. All plates were maintained in plastic containers with an open aperture to allow free exchange of gasses for the week-long duration of the experiments. The plastic chambers that housed the microplates contained moistened paper towels to inhibit plate evaporation. A Unisense oxygen microsensor was used to confirm that O_2_ was present and wells were well-mixed at depth in oxic treatments, including those that contained H_2_S.

### Image processing

Cells were imaged using an Olympus IX-81 inverted microscope equipped with a long working distance 40x objective (NA 0.6,WD 2.7-4.0), a 17.28 megapixel DP73 color camera. Images were collected using CellSens Dimension (Olympus, Japan) software under constant (manually-set) exposure and white balance settings. ImageJ was used to subtract the background and convert the image to an XYZ color profile. A 40x40 pixel region of interest representing an area of cytoplasm with a low density of sulfur globules was chosen for quantification of dye change in each cell. The average pixel intensity of this region was then measured in the luminance channel (Y). Reciprocal intensity was calculated using the approach described in (29) for quantifying chromogen intensity. Change in intensity relative to time zero was reported and a Student’s T-test was used to determine the likelihood that the imaged intensities were distinct from the mean of the controls by chance alone (Table 1). Copying the selection region in ImageJ from image-to-image in the time series for an individual cell ensured that the same area in the cell was measured at each time point.

**Table 1:** Mean positive change in the reciprocal intensity of luminance on Day 7 relative to time zero. Treatments that are highly statistically different (p < 0.005, Student’s t-test) from the mean of the four control samples are indicated in boldface type. Treatments that are marginally statistically different (p < 0.05, Student’s t-test) from the mean of the four control samples are indicated in underlined type. Negative values were used to calculate standard error, but are reported in the table as 0 to indicate no positive change.

| Substrate | *Thiomargarita* cells | Sheath control | Diatom control | Water controls |
| --- | --- | --- | --- | --- |
| **Oxic** |  |  |  |  |
| No donor added | 3.503±2.951 (n=8) | 0 | 0 | 0±0.214 (n=2) |
| Succinate | 3.383±2.123 (n=8) | 5.825 | 0.662 | 0±3.246 (n=2) |
| Thiosulfate | All cells collapsed (n=0) | 0 | 0 | 0±4.68 (n=2) |
| Acetate | 0±1.907 (n=8) | 2.652 | 0 | 0±1.829 (n=2) |
| **Citrate** | **13.009±2.108 (n=8)** | 3.993 | 2.878 | 0±2.421 (n=2) |
| Formate | All cells collapsed (n=0) | 0 | 5.672 | 0±0.243 (n=2) |
| **H<sub>2</sub>S** | **45.307±4.552 (n=8)** | 21.192 | 15.061 | 11.355±3.855(n=2) |
| **Hypoxic** |  |  |  |  |
| No donor added | 5.082±2.262 (n=8) | 0 | 0.838 | 2.973±0.577 (n=2) |
| **Succinate** | **24.742±2.081 (n=8)** | 0 | 3.102 | 0±0.438 (n=2) |
| Thiosulfate | All cells collapsed (n=0) | 5.151 | 0.744 | 0±0.293 (n=2) |
| <u>Acetate</u> | <u>44.131±12.107 (n=8)</u> | 9.431 | 7.652 | 0±0.643 (n=2) |
| <u>Citrate</u> | <u>30.562±4.289 (n=4)</u> | 1.579 | 0 | 0±0.422 (n=2) |
| Formate | 4.044±1.599 (n=8) | 3.411 | 4.665 | 0±0.17 (n=2) |
| <u>H<sub>2</sub>S</u> | <u>61.519±4.487 (n=6)</u> | 39.412 | 50.986 | 38.421±7.886 (n=2) |
| **Anoxic** |  |  |  |  |
| No donor added | 1.104±4.6 (n=8) | 1.938 | 0 | 0±0.581 (n=2) |
| **Succinate** | **50.732±1.274 (n=3)** | 13.507 | 3.852 | 0±5.616 (n=2) |
| **Thiosulfate** | **73.387±1.954 (n=8)** | 25.755 | 1.857 | 0±0.693 (n=2) |
| **Acetate** | **42.177±3.902 (n=8)** | 12.302 | 4.654 | 0.091±1.096 (n=2) |
| **Citrate** | **37.528±4.031 (n=8)** | 22.271 | 0 | 3.03±0.333 (n=2) |
| **Formate** | **42.234±6.21 (n=8)** | 2.583 | 13.464 | 0.432±0.114 (n=2) |
| **H<sub>2</sub>S** | **40.927±5.647 (n=8)** | 25.126 | 20.326 | 11.176±2.783 (n=2) |
| **H<sub>2</sub>** | **61.997± 5.125 (n=8)** | 23.555 | 19.556 | 0±0.49 (n=2) |

### Genome analysis

In order to compare our findings to genomic results, Beggiatoaceae genomes that were annotated by the IMG pipeline, were queried using tools available on IMG/ER (version 4.570 for this study)(30). The queried genomes are *Beggiatoa leptomitiformis* D-402 (GOLD analysis project Ga0111282), *Beggiatoa alba* B18LD (Ga0024935), *Candidatus* Maribeggiatoa sp. Orange Guaymas (Ga0010502), *Beggiatoa* sp. PS (Ga0027801), *Beggiatoa* sp. SS (Ga0027802), *Candidatus* Thiomargarita nelsonii Bud S10 (<u>Ga0097846</u>, <u>Ga0063879</u>), *Candidatus* Thiomargarita nelsonii Thio36 (<u>Ga0025452</u>) and *Thioploca ingrica* (<u>Ga0060138</u>). With the exception of the freshwater strains of *Thioploca (31)* and *Beggiatoa*, all genomes are incomplete genomes generated by multiple displacement amplification of individual cells (17, 18, 32, 33).

## RESULTS AND DISCUSSION

### Tetrazolium dye staining and the metabolism of lithotrophic substrates

*Thiomargarita* spp. cells exhibited apparent metabolic responses to a variety of substrates, as indicated by formazan staining that was localized within the cell (Figure 1), and that increased in intensity over the duration of the seven-day experiment (Figures 1A,B, Figs. 2-3). Maximum staining generally occurred within 4-6 days (Figs. 2-3).

**Figure 1:**
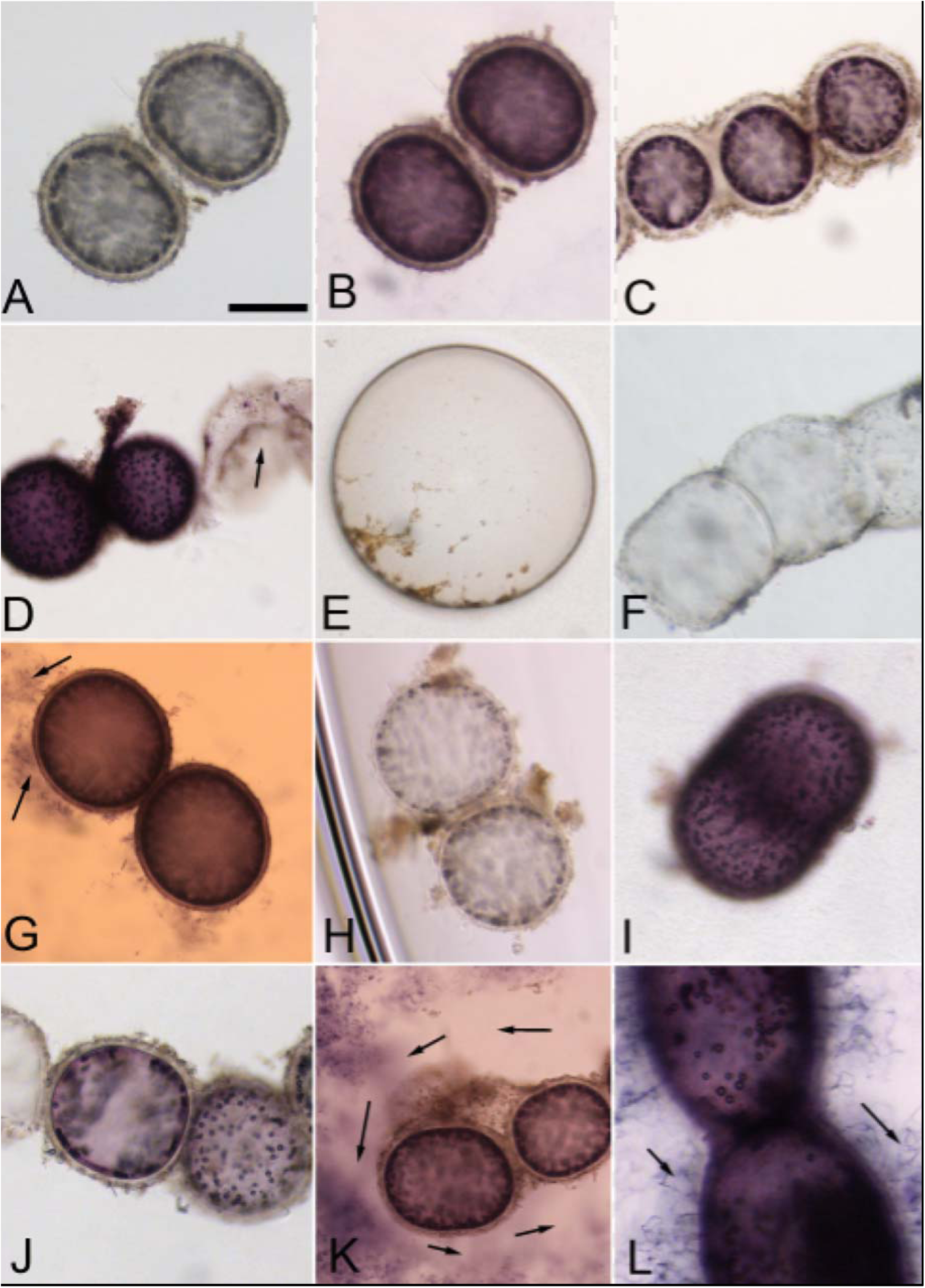
Cellular reductase activity reduces colorless tetrazolium to purple formazan product. Initially, *Thiomargarita* spp. cells incubated in the redox dye are colorless and the medium is light green in color. (**B**) Many intact cells under specific treatment conditions, presumably those that are metabolically active, stain a deep purple color and the extracellular medium changes to colorless or light pink, typically within two days. Panel B shows cells incubated under anoxic conditions in the presence of succinate. Staining of metabolically active cells was spatially separate and distinct from surrounding sheath material (**C).** Collapsed or damaged cells at the time of incubation did not exhibit a color change **(D**), showing exposure to acetate and hydrogen respectively, under anoxic conditions. Staining of metabolically active cells was distinct in intensity from control diatom frustules (**E**), and control sheath material (**F**). Under exposure to H_2_S, the extracellular medium assumed an orange hue that was readily distinguishable from the purple color change in the *Thiomargarita* spp. cells (**G**) and in biofilms of attached epibiont bacteria (arrows). Under anoxic conditions, cells and media showed the most extensive response, with very little color change observed with no additional electron donor added (**H**), and a strong staining response with the addition of other substrates such as H_2_ under anoxic conditions (**I, J**). In some cases, staining of extracellular bacteria were present as a diffuse stained cloud composed of small cells within the well. A zone characterized by an absence of staining and cells in the immediate vicinity of the *Thiomargarita* cells, suggests some sort of inhibition of these small bacteria (arrows in **K**). In other cases, such as under exposure to thiosulfate under anoxic conditions as shown here, stained filamentous epibionts could be observed anchored to the *Thiomargarita* cell/sheath (**L**). All images taken at 400x magnification, scale bar in A = ∼100 μm.

**Figure 2:**
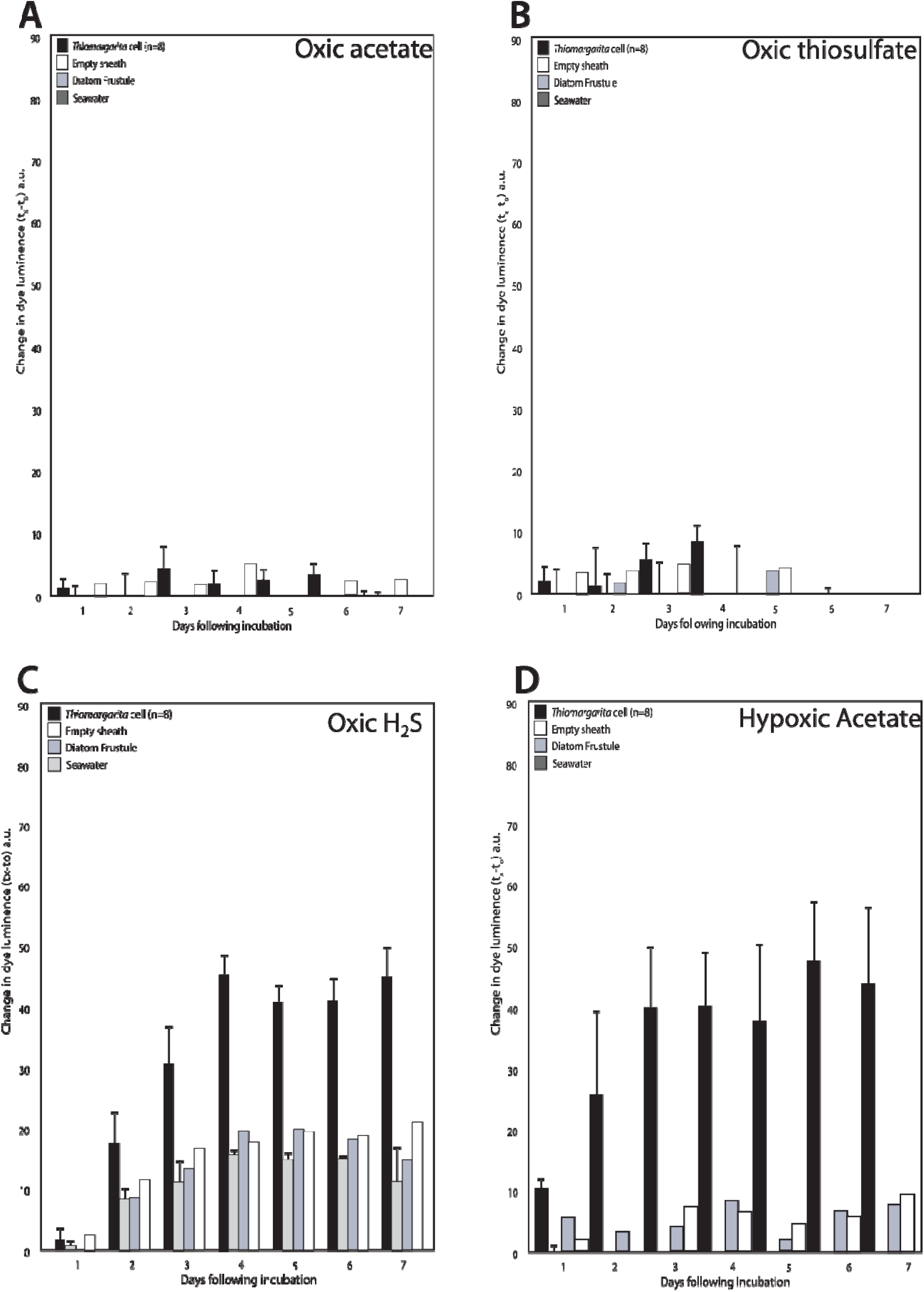
*Thiomargarita* cells exhibited very little staining response to acetate (**A**), thiosulfate (**B**), and other substrates, under oxygenated conditions relative to controls. However, H_2_S (**C**) and citrate (not shown), did induce a statistically-significant staining response under oxic conditions. Potentially significant staining was observed under hypoxic conditions in the presence of acetate (**D**), succinate, citrate, and H_2_S. Plotted here is the mean change in reciprocal intensity of luminance relative to day zero. Error bars indicate one standard error from the mean of the *Thiomargarita* incubations or controls.

Intense color changes were restricted to the *Thiomargarita* cell itself and did not extend to the sheath **(Figure 1B, C, J)**, except in cases where epibiont bacteria were stained as described below, or in the case of sulfide treatments, also described below. Collapsed or otherwise damaged cells at the time of incubation never showed a color change (Figure 1D), though some cells that did stain early in the experiment were sometimes observed to collapse in the later stages of the experiment. These observations suggest that formazan staining is a good indicator of initial cell viability, but that reductase activity alone was not sufficient to prevent eventual cell collapse in some specimens. Staining of controls, that included cell rinse water (not shown), diatom frustule (Figure 1E), and empty sheaths (Figure 1F) all exhibited very low intensities compared with *Thiomargarita* cell staining (Table 1; Figures 2-3).

Sulfur bacteria such as *Thiomargarita* are known for their ability to oxidize H_2_S using O_2_ or nitrate as an electron donor (3, 12). The intense tetrazolium dye staining of cells exposed to H_2_S under both oxic and anoxic conditions was highly statistically supported as being distinct from control staining (p < 0.005, Student’s t-test), while staining under sulfidic hypoxic conditions was marginally statistically distinct from control staining (p < 0.05, Student’s t-test) (Table 1). The genomes of *Thiomargarita* spp. contain genes for the oxidation of H_2_S via either a sulfide:quinone oxidoreductase (SQR), and/or flavocytochrome c (FCC) (17). In the treatments containing H_2_S, an orange color change was noted in the media (Fig. 1G). The orange color change, which only occurred in the presence of H_2_S, was distinguishable from the typical formazan purple color change that occurs within the cells (Fig. 1G). This orange coloration is likely the result of the abiotic reduction of tetrazolium by H_2_S. Despite the sulfide treatments having a higher background than other treatments, a dark purple color change in the cell could be differentiated from the orange background (Figure 1G, Figure 2C, Table 1).

Many sulfur-oxidizing bacteria are also known to be able to oxidize other sulfur-containing substrates such as thiosulfate. Thiosulfate addition to the experimental wells resulted in a increasingly strong formazan staining response under anoxic conditions (p < 0.005, Student’s t-test) (Figure 3D). However, significant staining was not observed with thiosulfate addition under oxic and hypoxic conditions (Table 1). Instead, cells were observed to collapse (Fig. 2B). All currently available genomes of Beggiatoaceae contain the genes for thiosulfate oxidation via a partial sox system (*soxABXY*) for the oxidation of thiosulfate to intracellularly stored sulfur granules. (17, 18) Thiosulfate is not as strong a reductant as H_2_S, which may have left *Thiomargarita* cells susceptible to oxidative stress under oxic and hypoxic conditions. Indeed, cell collapse was common with a variety of metabolic substrates under oxic conditions. Oxic treatments with H_2_S were the only exception.

**Figure 3:**
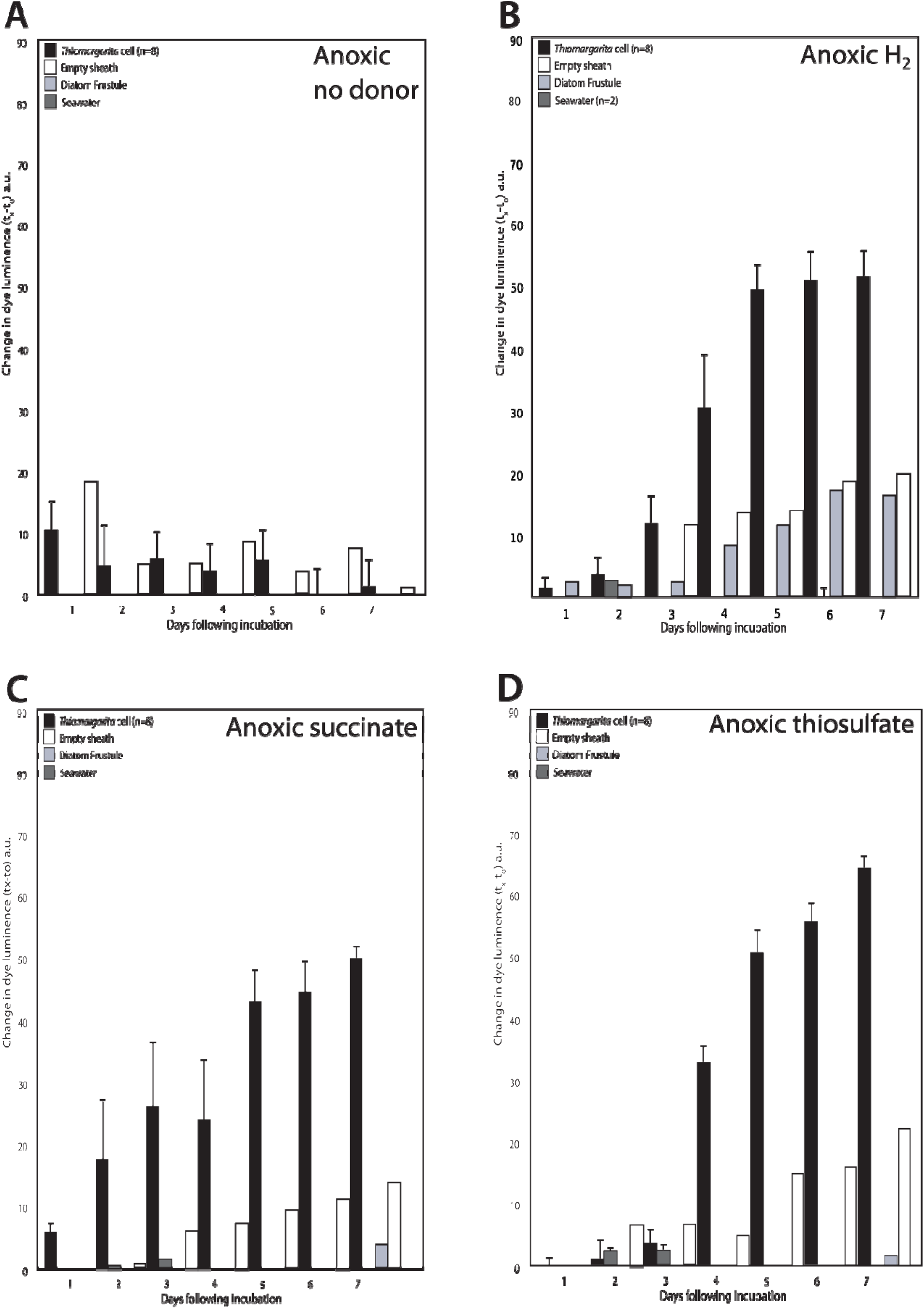
*Thiomargarita* cells and controls showed a weak response under anoxic conditions with the addition of a substrate that could not be differentiated from controls (**A**). With the addition of H_2_ (**B**), succinate (**C**), thiosulfate (**D**), H_2_S, formate, acetate, citrate, and formate (not shown), a statistically-significant staining response was observed under anoxic conditions. Plotted here is the mean change in reciprocal intensity of luminance relative to day zero. Error bars indicate one standard error from the mean of the *Thiomargarita* incubations or controls.

In addition to reduced sulfur compounds, some sulfur oxidizing bacteria are also known to use H_2_ as an electron donor for lithotrophic metabolism (34). Our staining results also showed high statistical support (p < 0.005, Student’s t-test) for formazan staining relative to controls when the cells were incubated in an anaerobic glove box containing 3% H_2_ (Figures 1D, 1I, 1J and 3B). Cells incubated without other supplied electron donors in an anoxic chamber without H_2_ showed no color change (Figures 1H, 3A). Recently, a chemolithotrophic strain of *Beggiatoa* sp., 35Flor, was found to use H_2_ as an electron donor under oxygenated conditions (35). 35Flor was not observed to oxidize H_2_ under anaerobic conditions, but *Beggiatoa* sp., 35Flor does not store nitrate to serve as an electron acceptor as *Thiomargarita* does. In addition to the microsensor studies that show H_2_ consumption by *Beggiatoa* sp. 35Flor, the genomes of *Ca.* T. nelsonii Thio36, and Bud S10 both contain genes for Ni-Fe hydrogenase (17, 18), as do those of other sulfur-oxidizing bacteria (36, 37). The source of H_2_ to *Thiomargarita* in nature is not clear, but fermentation by epibionts or other bacteria in the environment is a possible source.

### Organic acid metabolism

In addition to their canonical lithotrophic metabolism, some representatives of the Beggiatoaceae are known to metabolize organic acids (38). Organic acid degradation may be a common trait amongst sulfur-oxidizing gammaproteobacteria, since the generation of NADPH through the TCA cycle would coincide with the assimilation organic acids and would thus reduce the energetic costs of carbon fixation (39, 40). The genomes of *Ca.* T. nelsonii Bud S10, Thio36, and other members of the Beggiatoaceae, encode genes for a complete TCA cycle, including NADH dehydrogenase I (17, 18, 32, 33), and while succinate dehydrogenase is not electrochemically favored under most anaerobic conditions, it is tightly coupled with respiratory nitrate and nitrite reduction (41). Nitrate was provided in the medium (100 μM) and nitrate is also stored in the internal nitrate vacuole.

All the organic acids tested here yielded positive staining results under anoxic conditions, but staining results with organic acid exposure were more variable under hypoxic and aerobic conditions. We observed a positive staining response with the addition of citrate under oxic, hypoxic, and anoxic conditions, and succinate under hypoxic and anoxic conditions (Table 1). The utilization of exogenous organic acids requires specific inner membrane transporters. The genomes of a few Beggiatoaceae encode putative citrate transporters (Ga0060138_111857, BGP_2612) and a partial putative citrate transporter is in *Ca.* T. nelsonii Thio36 (Thi036DRAFT_00068870). Succinate is usually transported by proteins encoded by either one of two three-gene clusters, *dctABD*, where *dctA* is a permease and *dctBD* a two-component system for the activation of transcription (42) or similarly *kgtPSR* (43) or via a tripartite ATP-independent periplasmic transporter (TRAP), *dctPQM* (44). The two available *Thiomargarita* genomes contain a *dctB* gene but lack *dctAD*, although dctB is near the terminal end of a contig in the genome of Bud S10. While freshwater strains of *Beggiatoa* (*B. leptomitiforms* D-402 and *B. alba* B18LD) possess putative *dctPQM* genes, annotated genes for C4–dicarboxylate transporters are lacking in all marine strains. Thus, the capacity to transport succinate via known transporters in marine Beggiatoaceae appears to be lacking. However, the marine strains do possess a number of uncharacterized TRAP transporters that might serve as future targets for characterizing C4-dicarboxylate metabolisms. Acetate is another organic acid that is thought to be metabolized by certain sulfur bacteria. The microelectrode respiration experiments of Schulz and DeBeer showed the stabilization of oxygen/sulfide gradients around *Thiomargarita* when acetate was added to the experiments, although acetate itself was not found to steepen the oxygen gradient (16). Our formazan dye staining results show no statistical support for acetate use under oxic conditions, which is consistent with the results of Schulz and DeBeer. However, our results also show marginal statistical support for acetate metabolism under hypoxic conditions, and strong support for acetate use under anoxic conditions (Table 1, Figure 2D). Acetate can be metabolized by the tricarboxylic acid and glyoxylate cycles, and is preceded by the activation of acetate by acetate kinase and phosphotransacetylase (45) (known as the ack-pta pathway) and/or via an acetyl-CoA synthethase (known as the *acs* pathway). The ack-pta pathway occurs in the freshwater strains of the Beggiatoaceae, but not the marine strains. However, almost all Beggiatoaceae possess the acs pathway (18) and an acetate/cation symporter (e.g. Ga0063879_05139) (33). These genomic features are consistent with the ability of *Thiomargarita* spp. to take up and metabolize acetate from the environment, but our staining results suggest that this may only occur in *Thiomargarita* spp. under anoxic and perhaps hypoxic conditions.

Formate is both a waste product of fermentation and a potential electron donor that can be used for both aerobic and anaerobic respiration (46). The genomes of *Ca.* T. nelsonii Thio36 and *Beggiatoa* PS both contain genes that code for subunits of formate dehydrogenase. In addition to anaerobic respiration, energy-yielding reaction under anaerobic conditions is the formate-assisted cleavage of pyruvate via a pyruvate formate lyase enzyme. The genomes of both *T. nelsonii* contain three annotated pyruvate formate lyase genes and the gene occurs in most other Beggiatoaceae. The transport of exogenous formate could be mediated by a formate/nitrate transporter (focA), which is present in most marine strains of Beggiatoaceae, including both *T. nelsonii* genomes.

Although *Thiomargarita* spp. are thought to be more oxygen tolerant than other marine vacuolate sulfur bacteria, such as *Maribeggiatoa* and *Marithioploca* (16), the results presented here suggest that while *Thiomargarita* spp. may tolerate oxygen exposure, their metabolism(s) are most versatile under anoxic conditions. This is perhaps unsurprising given that *Thiomargarita* on the Namibian shelf are primarily found in sediments that are anoxic for much of the year (47). During the collection of samples used in our experiments, no oxygen was detectable in the lower water column as measured by Winkler titration, and in core-top waters as measured using a hand-held oxygen meter. Limited tolerance to O_2_ is also suggested by recent genomic results for most Beggiatoaceae. All Beggiatoaceae strains have cbb3-type cytochrome c oxidase and some also possess bd-type cytochromes, both of which are specific to hypoxic conditions. However, both *T. nelsonii* Thio36 and *Beggiatoa* PS possess the more oxygen-tolerant cytochrome c oxidase (*coxABC, cyoE*), which suggests greater tolerance in some strains of Beggiatoaceae. However, catalase, which is used to ameliorate oxidative stress, is not present in the genomes of marine Beggiatoaceae, occurring only in the freshwater strains. However, most Beggiatoaceae genomes contain a superoxide dismutase and cytochrome c peroxidase, while most marine strains also possess a desulfoferrodoxin. The enzymes coded for by these genes may provide protection against O_2_ and reactive oxygen species during periodic exposure to O_2_. Additionally, H_2_S, which can scavenge reactive oxygen species (48), may serve as an extracellular antioxidant under conditions in which sulfide is fluxing into oxygenated waters. Under anoxic conditions, oxidized forms of inorganic nitrogen whether exogenous, or stored within the vacuole, serve as terminal electron acceptors (3, 49). Both *Ca.* Thiomargarita nelsonii Bud S10 (17) and Thio36, possess a complete denitrification pathway to include both membrane bound (*nar*) and periplasmic (*nap*) nitrate reductases and the capacity to reduce nitrite to ammonium (*nirBD*). Thus, our staining results are consistent with the genomes and the canonical knowledge of nitrate being used as an electron acceptor by certain members of the Beggiatoaceae.

### Epibiont microbial cells

In our experiments, we undertook imaging of individual formazan- stained *Thiomargarita* cells in order to differentiate *Thiomargarita* metabolism from those of attached bacteria. These attached bacteria exhibited spatially-distinct staining from *Thiomargarita* under the microscope that couldn’t be differentiated with a typical microplate assay. Our microscope-based imaging approach had the added benefit of allowing us to observe the discrete staining of epibiont biofilms and filaments associated with *Thiomargarita* (Figure 1G, L). In some cases, stained filamentous epibionts could be observed anchored to the *Thiomargarita* cell/sheath (e.g. Figure 1L, arrows). In particular, dense accumulations of filamentous bacteria (Figure 1L) exhibited intense staining when exposed to H_2_ under anoxic conditions. While filamentous bacteria were observed in some control wells, and in zones of the well distal to *Thiomargarita,* their accumulations were far denser in the vicinity of *Thiomargarita* cells (Figure 1L). This observation suggests the possibility that *Thiomargarita* spp. cells are involved in syntrophic interactions with their epibionts. For example, *Thiomargarita* may supply attached sulfate-reducing bacteria with organic acids or sulfate, while the sulfate-reducing bacteria produce sulfide or other reduced sulfur intermediates that can be oxidized by *Thiomargarita (12)*. In these experiments H_2_ may have served as an electron donor for sulfate-reducing bacteria that then produced sulfide that was oxidized by the *Thiomargarita* cells. However, as discussed above, *Thiomargarita* spp. has the genetic potential to oxidize H_2_ directly.

We also observed under anoxic conditions in the presence of H_2_S, that small bacteria that stained purple were present throughout the medium, except in clear cell-free zones we observed immediately surrounding *Thiomargarita* cells and chains (Figure 1K, arrows). These zones may represent the inhibition of *Thiomargarita* on other small sulfur-oxidizing bacteria, perhaps through the drawdown of H_2_S in the vicinity of the *Thiomargarita* cells. Additional investigation, including alternative approaches such as gene expression studies, will be needed to further evaluate these interpretations.

### Conclusions

Our observations of reductase-mediated formazan staining in *Thiomargarita* cells exposed to a variety of organic and inorganic substrates are consistent with recent genomic results that suggest metabolic plasticity in *Ca.* Thiomargarita nelsonii. There are however, some limitations and caveats in drawing broad conclusions from our results. It is possible that the incubation media formulations or concentrations we used here, or other aspects of the experimental design, resulted in responses from *Thiomargarita* that do not represent their metabolic activities in nature. Yet the results presented here are broadly consistent with genomic data currently available for *Ca.* T. neslonii, and consistent with the sediment-hosted habitat of these bacteria that is anoxic for much of the year. We used both the spherical *Thiomargarita* cells that are typical of *Thiomargarita namibiensis* and the cylindrical forms that are more typical of Ca. *T. nelsonii* for this study (1). However, the large number of cells used in the experiment with the difficulty in amplifying *Thiomargarita’s* 16S rRNA gene due to multiple introns contained therein prevented a detailed phylogenetic characterization of the cells used for the assay (50). As such, we cannot say that our results apply broadly to the multiple candidate *Thiomargarita* species that can sometimes co-occur in sediments off Namibia (1, 51). While cultivation will ultimately be necessary for rigorous testing of the physiologies of *Thiomargarita* spp., for now, these results provide additional evidence in support of recent genomic and microsensor findings of metabolic versatility and/or mixotrophy in the genus *Thiomargarita* and suggest cultivation approaches that include both reduced sulfur compounds and organic substrates perhaps under extremely low oxygen conditions.

Tetrazolium/formazan redox dyes have long been used for studying bacteria, primarily in plate-based assays. Our expansion of the use of these dyes to the microscopic examination of individual stained cells, may be broadly useful for assaying metabolism in mixed communities and other uncultivated microorganisms, and for validating genome-derived assessments of physiological capabilities.

## FUNDING INFORMATION

This work was supported by a grant from the Simons Foundation (341838 J.B.), by an Alfred P. Sloan Research Fellowship (BR2014-048), and by a grant from the National Science Foundation (EAR-1057119), and by the Regional Graduate Network in Oceanography Discovery Camp that is itself funded by the Agouron Institute and the Scientific Committee for Oceanographic Research (SCOR).

### ACKNOWLEDGEMENTS

We gratefully acknowledge the assistance of the staff and students of the RGNO Discovery Camp, the University of Namibia, the Namibian Ministry of Fisheries, Kurt Hanselmann, Daniel Montlucon, Deon Louw, Richard Horaeb, Verena Carvalho, Bronwen Currie, the crew of the R/V *Mirabilis*.

